# HIV cell-to-cell spread slows evolution of drug resistance

**DOI:** 10.1101/2020.09.04.283192

**Authors:** Jessica Hunter, Sandile Cele, Laurelle Jackson, Jennifer Giandhari, Tulio de Oliveira, Gila Lustig, Alex Sigal

## Abstract

Many enveloped viruses such as HIV have evolved to transmit by two infection modes: cell-free infection and cell-to-cell spread. Cell-to-cell spread is highly efficient as it involves directed viral transmission from the infected to the uninfected cell. In contrast, cell-free infection relies on chance encounters between the virion and cell. Despite the higher efficiency of cell-to-cell spread, there is substantial transmission by cell-free infection in conjunction with cell-to-cell spread. A possible reason is that cell-free infection offers a selective advantage by increasing sensitivity to factors interfering with infection, hence accelerating evolution of resistance relative to cell-to-cell spread alone. Here we investigated whether a combination of cell-free infection and cell-to-cell spread confers a selective advantage in experimental evolution to an antiretroviral drug. We maintained HIV infection using coculture of infected with uninfected cells in the face of moderate inhibition by the reverse transcriptase inhibitor efavirenz. We tested the effect on the rate of drug resistance evolution of replacing one coculture infection cycle with an infection cycle involving cell-free infection only, and observed earlier evolution of drug resistance mutations to efavirenz. When we increased selective pressure by adding a second reverse transcriptase inhibitor, emtricitabine, infection with the cell-free step consistently evolved multidrug resistance to both drugs and was able to replicate. In contrast, infection without a cell-free step mostly failed to evolve multidrug resistance. Therefore, HIV cell-to-cell spread decreases the ability of HIV to rapidly evolve resistance to inhibitors, which is conferred by cell-free infection.

**Author summary:** Cell-to-cell spread of HIV differs from cell-free, diffusion-based HIV infection in that viral transmission is directed from the infected to the uninfected cell through cellular interactions. Cell-to-cell spread has been recognized as a highly efficient infection mode that is able to surmount inhibition by antibodies and antiretroviral drugs. However, the effect of HIV cell-to-cell spread on the rate of evolution of viral resistance to infection inhibitors has not been studied. Here we used experimental evolution to investigate the effect of cell-to-cell spread versus cell-free infection on the emergence of drug resistance mutations to one or a combination of antiretroviral drugs. We found that replacing one infection cycle in experimental evolution with cell-free infection, where the filtered supernatant from infected cells, but not the cellular fraction, is used as the viral source, results in more rapid evolution of resistance. The consequences are that multidrug resistance consistently evolves with a cell-free viral cycle, but not when infection is solely by coculture of infected and uninfected cells. A possible consequence is that in environments where HIV cell-to-cell spread may predominate and some residual viral replication occurs in the face of ART, the emergence of drug resistance mutations would be delayed.

## Introduction

Despite the overall success of antiretroviral therapy (ART) at suppressing HIV replication, evolution of drug resistance remains a considerable concern as it leads to the replication of HIV in the face of ART due to acquisition of drug resistance mutations. Incomplete viral suppression of HIV because of treatment interruptions (1–7), and lowered drug levels in some anatomical compartments (8–12) contribute to the selection of drug resistance mutations (13, 14). Drug resistance mutations enable HIV to replicate in the face of what should be suppressive ART concentrations if sufficient mutations are accumulated and result in multidrug resistance (10, 15). The prevalence of drug resistance mutations in HIV infected individuals on ART is about 10% (16, 17), (see also the WHO HIV Drug Resistance Report 2019 at https://www.who.int/hiv/pub/drugresistance/hivdr-report-2019/en/).

A mechanism for drug insensitivity distinct from acquisition of drug resistance mutations is HIV cell-to-cell spread (18–22). Cell-to-cell spread involves the directed transmission of virions from one cell to another at close range, generally through a virological synapse or other structure which minimizes the distance HIV has to diffuse to reach the uninfected cell (23–30) but does not involve fusion (31). It has been shown to be an efficient mode of infection which decreases sensitivity to antiretroviral drugs (18, 19, 32, 33) and neutralizing antibodies (34–37). It also allows infection under unfavourable conditions (36). This includes infection of cells such as resting T cells and macrophages which have low numbers of CD4 entry receptors (38, 39). In addition to increased efficiency, HIV cell-to-cell spread makes infection more rapid (22) and can more easily cross mucosal barriers (40).

Despite the advantages to the virus of infecting by cell-to-cell spread, a considerable amount of virus is not localized to cell-to-cell contacts (25) and transmits by cell-free infection (41–43). More generally, many enveloped viruses are able to infect by both cell-to-cell and cell-free modes (44). This may occur because cell-to-cell spread can be cytotoxic (33, 45) or cell-free virus is needed for transmission between individuals. Here we investigated the possibility that cell-free HIV infection, due to its increased sensitivity to inhibitors, could confer a selective advantage by reducing the time to evolution of drug resistance (8, 46, 47).

We performed *in vitro* evolution in the face of the antiretroviral drug efavirenz (EFV) using coculture of infected with uninfected cells. We used a concentration of EFV similar to that predicted upon treatment interruption of several days, due to the increased half-life of EFV relative to the other drug components of the ART regimen (7, 48–53). We introduced a cell-free infection cycle, where we harvested the cell-free virus by filtering out the infected cells and used it as the sole infection source for one infection cycle. The other infection cycles occurring by coculture of infected and uninfected cells which allows cell-to-cell spread (18). We observed a faster fixation of an EFV drug resistance mutant with the cell-free infection cycle relative to coculture alone. Upon transfer of infection with the cell-free step to a culture containing a two drug combination of EFV and the reverse transcriptase inhibitor emtricitabine (FTC), infection containing a cell-free infection cycle was able to evolve drug resistance to both drugs, while infection without the cell-free step failed to evolve multidrug resistance. Therefore, cell-free infection confers the ability to rapidly evolve to selective pressure.

## Results

### Faster evolution of drug resistance to EFV with cell-free infection

We reasoned that in cell-to-cell spread, which occurs when infected (donor) cells and uninfected (target) cells are cocultured, the frequency of a drug resistant mutant would rise gradually in the face of moderate drug pressure. This would be because the mutant needs to supplant the still replicating drug sensitive (wild-type) HIV genotype. In contrast, cell-free infection is more sensitive to inhibitors, and so wild-type virus would be effectively cleared even at moderate levels of drug.This would rapidly increase the frequency of the drug resistant mutant (Figure 1A).

**Fig. 1.**
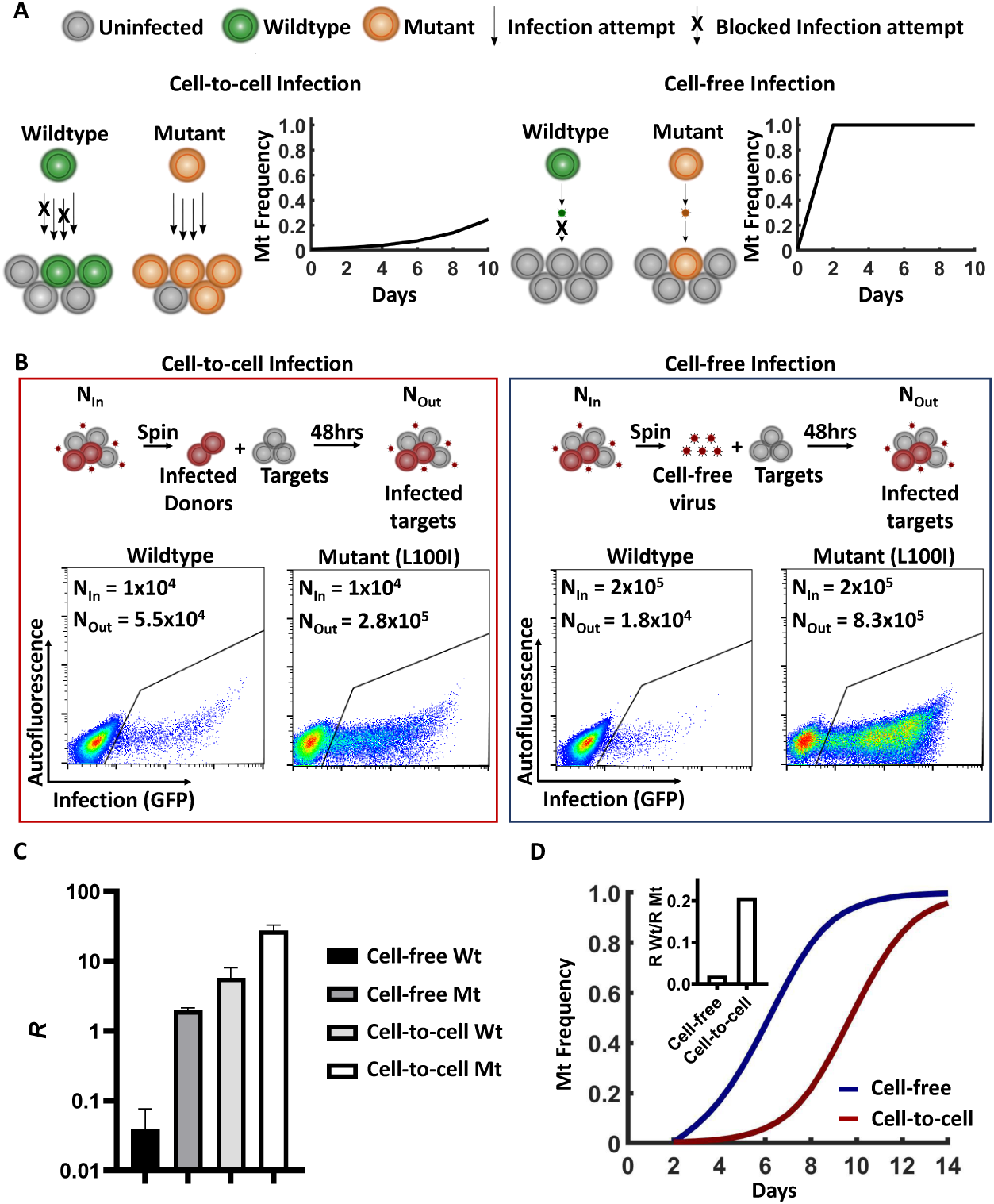
Wild-type virus is predicted to be more rapidly supplanted by the drug resistant mutant with cell-free infection relative to cell-to-cell spread. A) Schematic. Orange cells are mutant infected, green are wild-type infected, grey uninfected. Arrows represent infection attempts, with an “x” denoting an infection attempt blocked by drug. B) Measurement of the replication ratio (*R*), defined as the number of infected cells at the end of 48 hours of infection (*N*_*out*_) divided by input number of infected cells added directly or from which cell-free virus was harvested at infection start (*N*_*in*_). The cellular and cell-free fractions were derived by spinning the sample and using the infected cells from the pellet (cell-to-cell, left plot) or the filtered supernatant of the same infected cells (cell-free, right plot) for infection. Infected cells are in the GFP positive gate. Left plots shows *N*_*out*_ per mL for wild-type and mutant infection when infection was by cell-to-cell spread and *N*_*in*_ per mL was 1 × 10^4^. Right plots shows *N*_*out*_ per mL for wild-type and mutant infection when infection was by cell-free virus and *N*_*in*_ per mL was 2 × 10^5^. C) Measured *R* values for wild-type and L100I mutant HIV for infection by cell-free and cell-to-cell. D) Expected frequency of the drug resistant mutant with (blue line) and without (red line) a single cycle of cell-free infection. Frequency was calculated as 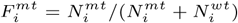, where 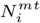 and 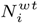 are the number of mutant and wild-type infected cells respectively at infection cycle *i*, and where *N*_*i*_ = *RN*_*i*−1_. Inset shows the ratio of wild-type to mutant *R* values for cell-free infection and cell-to-cell spread.

To experimentally measure the effect of drug on wild-type and mutant virus in cell-free infection and cell-to-cell spread, we determined the replication ratio for both infection modes in the face of 20nM of EFV. The replication ratio with cell-to-cell spread (*R*_*cc*_) was derived by measuring the number of infected cells (*N*_*out*_) 2 days (approximately 1 viral cycle) post-infection divided by the number of input infected cells (*N*_*in*_). The cell-free replication ratio (*R*_*cf*_) was derived as for cell-to-cell spread, except that only the supernatant from *N*_*in*_ was used (Figure 1B). To determine *R*_*cf*_, input was supernatant from 2 × 10^5^ cells/mL infected with wild-type or the L100I EFV resistance mutant, where the L100I confers about 10-fold resistance to EFV (https://hivdb.stanford.edu/cgi-bin/PositionPhenoSummary.cgi). We obtained a mean *N*_*out*_ of 1.8 × 10^4^ ± 7.9 × 10^3^ infected cells for the cell-free infection with wild-type virus. When L100I mutant virus was used, the mean *N*_*out*_ increased to 8.3 × 10^5^ ± 1.8 × 10^4^. To measure *R*_*cc*_, we used a lower input of 1 × 10^4^/mL of cells infected with wild-type or the L100I EFV resistance mutant to prevent saturation of infection. We obtained a mean *N*_*out*_ of 5.5 × 104 ± 1.7 × 10^4^ for infection with wild-type and 2.8 × 105 ± 5.4 × 10^4^ for infection with the L100I mutant virus respectively. The mean *R*_*cf*_ was therefore 0.039 ± 0.032 for wild-type virus and 2.0 ± 0.16 for the L100I mutant (Figure 1C). The mean *R*_*cc*_ was 5.5 ± 1.7 for wild-type virus and 28 ± 5.4 for the L100I mutant. Therefore, the effect of 20nM of EFV on coculture infection was moderate: wild-type infection could still replicate and expand, as *R >* 1. In contrast, the effect of EFV on cell-free infection of wild-type was much stronger. *R*_*cc*_ of the L100I mutant was approximately 5-fold higher compared to wild-type. In contrast, the L100I mutant increased *R*_*cf*_ approximately 50-fold.

To determine the expected effect of this difference, we calculated the expected frequency of the drug resistant mutant in the total viral pool over time (Figure 1D, Materials and Methods) starting from the measured frequency of the L100I mutant (SFig. 1). We either included or excluded one cell-free infection step at the first infection cycle. The calculation showed that with an initial cell-free infection cycle, the frequency of the L100I EFV resistant mutant was substantially higher at the initial timepoints relative to cell-to-cell spread alone. At later timepoints post-infection, the L100I mutant supplanted the wild-type whether or not a cell-free step was included.

To examine whether a cell-free infection step could accelerate evolution of drug resistance, we performed *in vitro* evolution experiments in the face of 20nM EFV by the coculture of infected with uninfected cells. Two cycles of infection were first performed in the absence of drug to obtain a quasispecies (Materials and Methods), allowing selection from a pre-existing pool of single drug resistant mutations which were present at a low but detectable frequency (SFig. 1). Ongoing infection was maintained in the presence of drug by the addition of new uninfected cells every 2 days (Figure 2A). We included a cell-free infection cycle by pelleting the infected cells and filtering the supernatant to obtain cell-free virus and infected cells from the same infected cell population. We then infected new cells by using either the entire cellular fraction or the entire filtered supernatant from the cellular fraction during the first infection cycle in the presence of EFV (Figure 2A). In the latter case, infection would be exclusively by the cell-free mode. 20nM EFV monotherapy, the drug concentration used, may occur in individuals on EFV based ART regimen after several days of treatment interruption due to the longer half-life of EFV relative to other common ART components (7, 48–53).

**Fig. 2.**
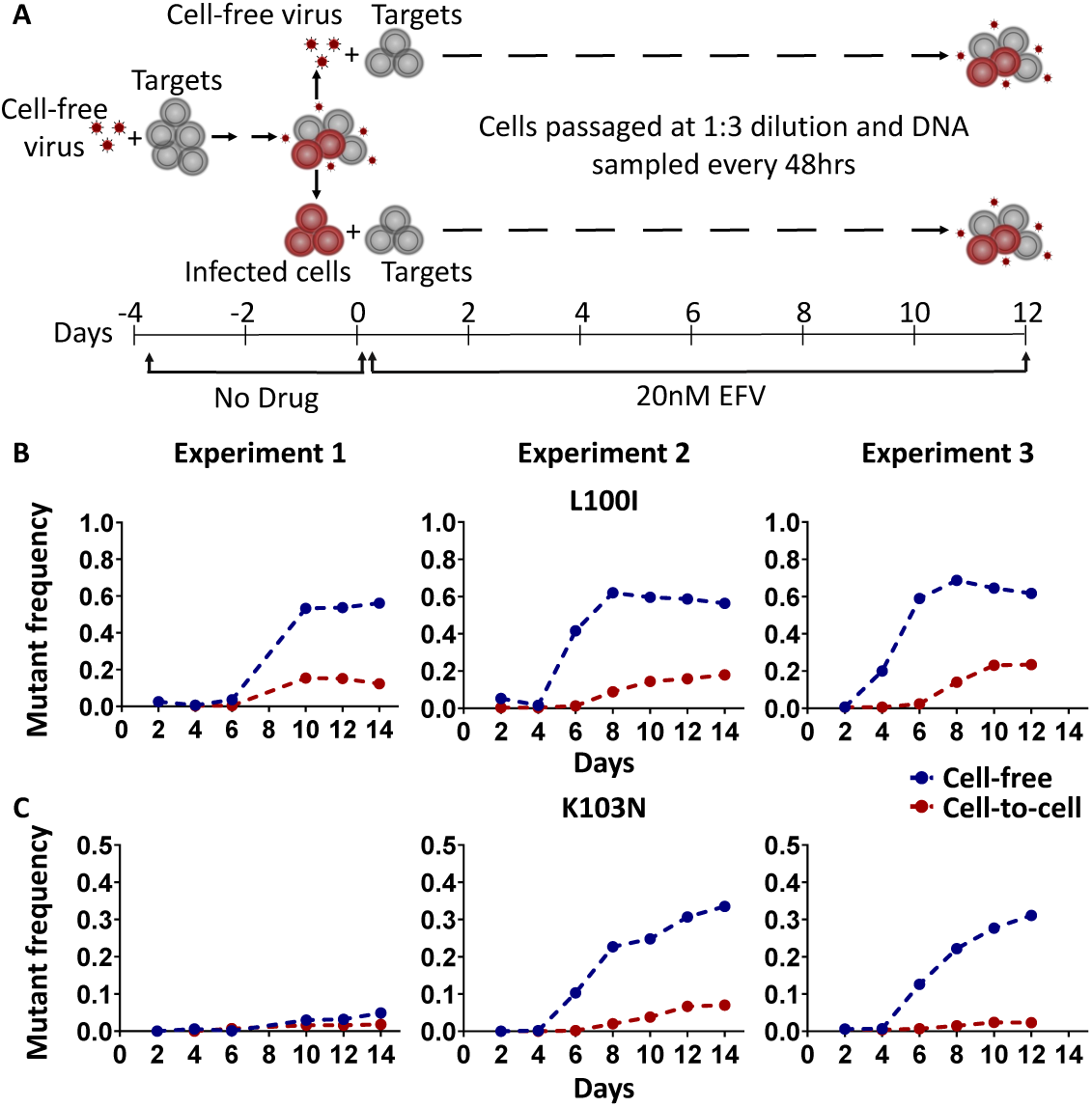
Frequencies of L100I and K103N EFV resistance mutations rise more rapidly with a cell-free infection step. A) Schematic of the experimental procedure. Targets denote uninfected cells to which cell-free virus or infected cells are added. B) Frequencies of the L100I EFV resistance mutation with a cell-free step (blue points) and without a cell-free step (red points) in three independent experiments. C) Frequencies of the K103N EFV resistance mutation with and without a cell-free step in the same three independent experiments as (B).

Upon passaging of infection with 20nM EFV, we obtained evolution of the L100I drug resistance mutation (Figure 2B), and with a delay the K103N drug resistance mutation in two out of three experiments (Figure 2C). The K103N is the more commonly detected EFV resistance mutation in the clinical setting and confers approximately 20-fold resistance to EFV (https://hivdb.stanford.edu/cgi-bin/PositionPhenoSummary.cgi).

In all experiments, a cell-free infection cycle led to a more rapid increase in and higher final frequencies of the L100I mutation. By day 12 post-infection, mean mutant frequency was 0.58 ± 0.040 for evolution with a cell-free infection cycle, and 0.11 ± 0.076 without the cell-free infection cycle.

The increase in frequency of the K103N mutation was also accelerated for infection with a cell-free infection cycle in experiments where the K103N evolved to substantial levels. Mean K103N mutant frequency at day 12 post-infection was 0.21 ± 0.15 in the infection with a cell-free step, and 0.035 ± 0.027 when infection was in the absence of the cell-free step. Therefore, adding a cell-free infection cycle increased the frequencies of both drug resistance mutations in the experiments where they evolved.

### Cell-free infection but not cell-to-cell spread leads to the evolution of multidrug resistance

To determine whether infection with a cell-free step under EFV alone conferred a selective advantage to HIV with a combination of drugs, we added a second drug (FTC) after a period of monotherapy with EFV. As before, the supernatant containing cell-free virus was separated and used to establish an infection with a cell-free step in the face of EFV. The cellular fraction was used for infection without a cell-free infection cycle in the face of EFV. Evolution was carried out at 20nM EFV for the first 3 viral cycles (6 days). Thereafter, a drug combination of 20nM EFV and 770nM FTC was used, where the FTC concentration used similar to that found in study participants on ART (48, 54, 55). Infected cells were transferred from the one drug to the two drug regimen so that in the two drug regimen infection was initiated with 1% infected cells.

We performed three evolution experiments where we tracked the frequency of drug resistance mutations through time (Figure 3). We observed an increase in the frequency of multidrug resistance (mutations to both EFV and FTC) in the presence of the cell-free step: The EFV resistance mutations L100I or K103N were linked to FTC mutations M184V or M184I. Both M184V and M184I confer high level resistance to FTC (https://hivdb.stanford.edu/cgi-bin/PositionPhenoSummary.cgi).

**Fig. 3.**
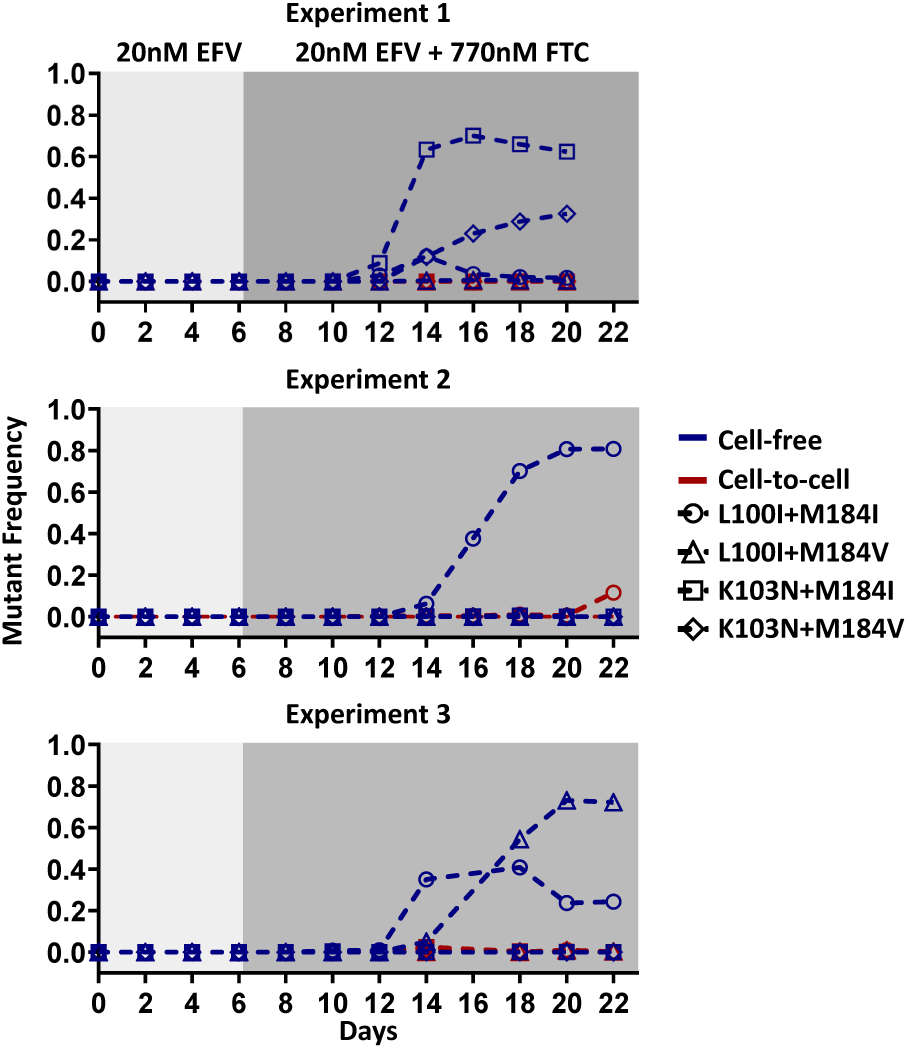
HIV infection with a cell-free infection cycle evolves multidrug resistance. Frequencies of drug resistant mutations for infection with a cell-free infection cycle (blue) and without a cell-free infection cycle (red) for three independent evolution experiments. light grey shading indicates the days at which the infection was carried out under EFV monotherapy conditions (20nM EFV). Dark grey background shading indicates the days at which the infection was carried out in the face of the two drugs (20nM EFV and 770nM FTC).

In the first experiment, the infection with a cell-free step evolved the L100I single mutant by day 6 under EFV monotherapy (SFig. 2A). This mutation was supplanted by the K103N single mutant and the L100I/M184I, K103N/M184I, and K103N/M184V multidrug resistant mutants during evolution in the face of EFV and FTC. The K103N and the L100I M184I variants were in turn out-competed, with the K103N/M184I and K103N/M184V multidrug resistant mutants dominating by day 16 (Figure 3, blue squares and diamonds, respectively). In the second experiment, the infection with a cell-free step showed detectable L100I at day 6 (SFig. 2A). The L100I/M184I arose to detectable levels by day 14 and dominated infection by day 18 of the experiment (Figure 3, blue circles). In the third experiment, by day 6 the infection with a cell-free step evolved the L100I mutation to a frequency of 0.4 (SFig. 2A). This mutant was supplanted by the L100I/M184I and L100I/M184V multidrug resistance mutants in the presence of the combined regimen of EFV and FTC (Figure 3, blue circles and triangles, respectively). Infection without a cell-free step failed to evolve multidrug resistance (Figure 3). The only possible exception was the L100I M184I resistance mutant, which started to be detectable on day 22 of experiment 2 (Figure 3, red circles).

### Increased drug pressure leads to more rapid evolution of drug resistance in the absence of cell-free infection

The slower evolution of drug resistant mutations without a cell-free step is consistent with the drug exerting weaker selective pressure in cell-to-cell spread compared with cell-free infection (18, 33). To test the result of increasing selective pressure, we performed the evolution experiments using a higher EFV concentration of 40nM. As previously, the cell-free and cellular fractions were separated from the same infected cell population. The infected cells were used to establish an infection without a cell-free step. After passaging the infection at 40nM EFV, there was an increase in both the rate of evolution as well as the maximal frequency of the L100I mutation (Figure 4A) and the K103N mutation (Figure 4B) when compared to 20nM EFV. Mean mutant frequency for the L100I mutant at day 8 was 0.43 ± 0.15 for infection in the face of 40nM EFV, close to its maximum value in these experiments. In contrast, the L100I frequency was 0.11 ± 0.036 at 20nM EFV. Similarly, the K103N mutant reached its maximum frequency by day 8 at 40nM. The mean K103N frequency at day 8 at 20nM was 0.094 ± 0.070. In contrast, the frequency was 0.018 ± 0.0035 at 20nM EFV. Therefore, addition of selective pressure by increasing drug concentration results in more rapid evolution by cell-to-cell spread.

**Fig. 4.**
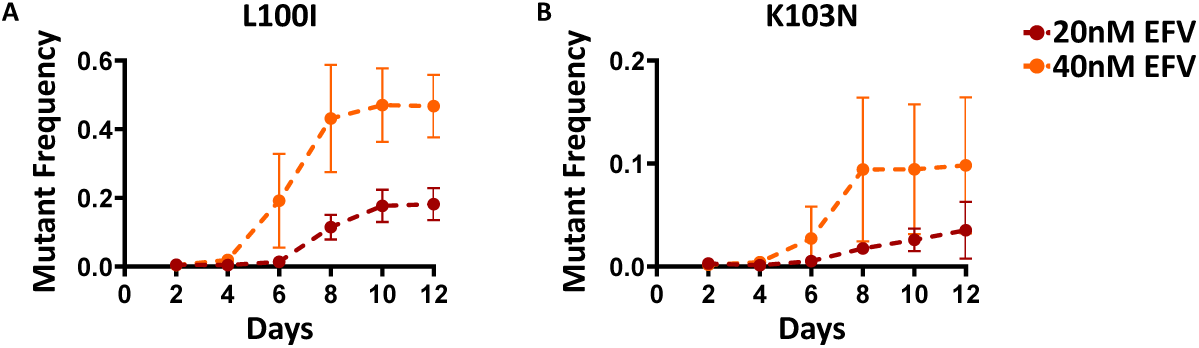
Increasing drug concentration in coculture infection increases the rate of drug resistance evolution. A) L100I and B) K103N mutation frequencies for infection without a cell-free infection cycle passaged either at 20nM EFV (red) or at 40nM EFV (orange). Mean and standard deviation from 3 independent experiments.

### Increased drug pressure leads to a minor increase in drug resistance frequency with cell-free infection

We asked whether the evolution of drug resistant mutations with a cell-free step would be further accelerated if drug pressure was increased. We therefore performed the experiments using 40nM EFV, as above, except that the cell-free fraction was used to establish the infection in the face of drug. Interestingly, when we compared the result to the same experiment carried out at 20nM EFV, there was a minor difference in the time to maximal frequency of the L100I mutation (Figure 5A). The L100I mutation frequency reached the near maximal mutation frequency on day 6 for 40nM EFV and day 8 for 20nM EFV. The maximal difference in frequency occurred on day 6, with mean frequency being 0.62 ± 0.080 at 40nM EFV and 0.35 ± 0.28 at 20nM EFV. The rate and the maximal frequency of the K103N mutation for the infection with a cell-free step was moderately increased at 40nM EFV compared to 20nM EFV. (Figure 5B). At the last timepoint tested (day 12), mean frequency for K103N was 0.41 ± 0.20 at 40nM EFV, compared to 0.22 ± 0.16 at 20nM EFV. Therefore, while higher EFV led to an increase in the rate of selection of K103N, 20nM EFV was close to the maximal selective pressure required for rapid evolution of L100I.

**Fig. 5.**
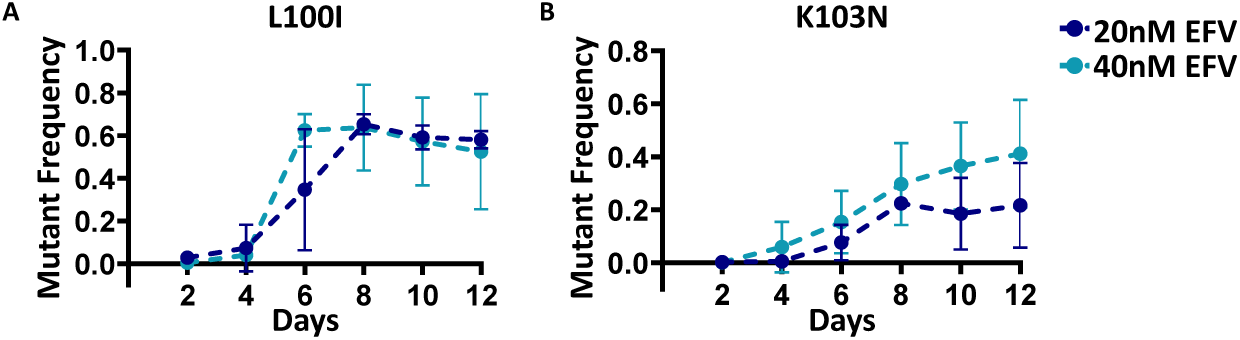
Frequency of resistance mutations with a cell-free infection cycle under increased selective pressure. A) L100I and B) K103N mutation frequencies for infection with a cell-free step passaged either at 20nM EFV (dark blue) or at 40nM EFV (light blue). Mean and standard deviation from 3 independent experiments.

## Discussion

We demonstrated that evolution of drug resistance occurs more rapidly with a cell-free infection step. The ability to rapidly adapt to changing environmental conditions may be one explanation why HIV is transmitted between cells by cell-free infection in conjunction with cell-to-cell spread, despite the greater efficiency of the latter (19, 41, 56, 57). In the experiments, we removed all cell-to-cell spread for one infection cycle by infecting only with the cell-free virus in the filtered culture supernatant. Even though an infected cell can transmit HIV by both cell-free infection and cell-to-cell spread, cells infected at a distance from the virus producing cell, where contact between the infected and uninfected cell does not occur, may be infected by cell-free infection alone. The consequences for evolution should be captured by our experiments.

We observed that the L100I mutation was the first mutation to arise, and the rate at which it dominated the viral population was further accelerated by a cell-free infection step. The K103N mutation showed a slower increase. Yet, with both mutations, a single cycle of cell-free infection accelerated the rate at which they became predominant. Therefore, accelerated evolution of drug resistance with cell-free infection is not mutation specific. The effect of s cell-free infection cycle is similar to increasing the selective pressure in coculture infection (Figure 4). The moderate selective pressure achieved with 20nM EFV was close to the maximal level needed to select for drug resistance mutants with a cell-free infection step (Figure 5). The experimentally observed effects of a cell-free step (Figure 2) were stronger than those predicted by the higher selective pressure alone (Figure 1D). Additional processes may therefore be involved which decrease the rate at which the drug resistant mutant out-competes the wild-type in cell-to-cell spread, such as interactions between the mutant and wild-type through a process of complementation (20).

The ability to rapidly evolve drug resistance may be particularly important when evolving resistance to a regimen containing multiple antiretroviral drugs, and the primary reason current regimens contain multiple antiretroviral compounds is to prevent the evolution of drug resistance (8), as even single antiretrovirals may be sufficiently potent in suppressing HIV (58). When we introduced FTC as the second drug after EFV monotherapy, only infections with a cell-free step evolved resistance to both drugs, with the exception of one experiment, where multidrug resistance became detectable in the last timepoint of infection in the absence of a cell-free infection cycle. EFV monotherapy occurs in the clinical setting during treatment interruptions, as EFV has a longer half-life relative to FTC and the reverse transcriptase inhibitor tenofovir which are usually co-formulated with it (2, 48, 50–53). Monotherapy provides the opportunity for HIV to accumulate drug resistance mutations in a stepwise fashion (10, 15). If HIV evolves more rapidly by cell-free infection relative to cell-to-cell spread, infection by cell-free virus would allow it to quickly evolve drug resistance to the single drug during a window of monotherapy. As the other drugs are reintroduced, the replication of partially resistant virus should enable it to evolve additional drug resistance mutations. Conversely, microenvironments which favour cell-to-cell spread such as lymph nodes (59, 60) would not be expected to rapidly evolve drug resistance. If residual and compartmentalized HIV replication (12, 60, 61) does occur in some anatomical compartments, it may show a delay in drug resistance evolution. This may explain why modest non-adherence does not necessarily lead to failure of the ART regimen due to evolution of drug resistance (62–64). This is despite the expectation that, during such periods, HIV from the latent reservoir (65–72) could initiate cycles of viral replication.

By using both the cell-free and cell-to-cell infection modes, HIV is able to both rapidly evolve resistance to factors interfering with infection, and take advantage of an efficient infection mode once adaptation occurred. This makes the virus particularly suitable to survive long-term under conditions of an evolving immune response or partial drug inhibition where rapid evolution is key.

## Materials and Methods

### Inhibitors, viruses and cell lines

The antiretrovirals EFV and FTC were obtained through the AIDS Research and Reference Reagent Program, National Institute of Allergy and Infectious Diseases, National Institutes of Health. HIV molecular clone pNL4-3 was obtained from M. Martin. Viral stocks of NL43 were produced by transfection of HEK293 cells with the molecular clone plasmid using TransIT-LT1 (Mirus) transfection reagent. Supernatant containing released virus was harvested two days post-transfection and filtered through a 0.45 micron filter (GVS). The supernatant containing virus was stored in 0.5ml aliquots at −80°C. The L100I pNL4-3 molecular clone was generated as previously described (33). RevCEM cells were obtained from Y. Wu and J. Marsh. RevCEM-E7 cells were generated as described in (22). Briefly, the E7 clone was generated by subcloning RevCEM cells at single cell density. Surviving clones were subdivided into replicate plates. One of the plates was screened for the fraction of GFP expressing cells upon HIV infection using microscopy, and the clone with the highest fraction of GFP positive cells was selected. Cells were cultured in complete RPMI 1640 supplemented with L-Glutamine, sodium pyruvate, HEPES, non-essential amino acids (Lonza), and 10% heat-inactivated FBS (Hyclone).

### Measurement of replication ratios for cell-to-cell spread and cell-free infection

To calculate the replication ratio for cell-free infection (*R*_*cf*_), the supernatant of 2 × 10^5^ infected RevCEM-E7 cells were used as the input to 10^6^ cells/ml uninfected target RevCEM-E7 cells. For the calculation of the cell-to-cell replication ratio (*R*_*cc*_), 1 × 10^4^ infected donor RevCEM-E7 cells were added as the input to 10^6^ cells/ml uninfected target cells. After 48 hours of incubation at 37°*C*, the number of output infected cells determined by flow cytometry as the number of GFP expressing cells.

### Calculation of mutant frequency with and without a cell-free infection step

The calculation was performed using a Matlab 2019a script where mutant frequency was calculated as:

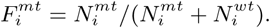

Here 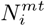 and 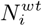 are the number of mutant and wild-type infected cells at infection cycle *i*, where each infection cycle is approximately 2 days.

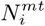 and 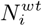 are determined as:

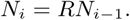

Here *R* is the replication ratio of the infection for wild-type or mutant virus using the cell-free infection (*R*_*cf*_) or cell-to-cell spread (*R*_*cc*_), and *N*_*i*−1_ is the number of wild-type or mutant infected cells in the previous infection cycle.

If a cell-free step was included, the first infection cycle uses 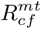 and 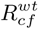 and the number of infected cells for mutant infection is calculated as:

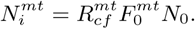

Similarly for wild-type virus,

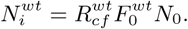

Here 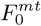 is the measured initial mutant frequency, where the frequency for the L100I mutant was used (*µ* = 5 × 10^−3^, *σ* = 5 × 10^−3^) and 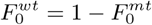. *N*_0_ is the initial number of infected cells and can be arbitrary.

If a cell-free step was excluded, and for all viral cycles after *i* = 1, the calculation uses 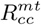 and 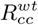, and the number of infected cells for mutant infection is calculated as:

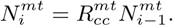

Similarly for wild-type virus,

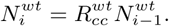

### Evolution Experiments

Two rounds of infection were performed in the absence of drug to establish a quasispecies, followed by multiple rounds of infection in the presence of EFV or EFV and FTC. The first round of infection in the absence of drug was initiated by cell-free infection of 2.5 × 10^6^ RevCEM-E7 cells with 5 × 10^8^ viral copies of HIV NL4-3 and incubated for 48 hours. In the second round of infection, cells infected in the first round were added to 3 × 10^6^ uninfected RevCEM-E7 cells to a final concentration of 0.5% infected cells and incubated for 48 hours. The infection was then separated into infected cells and cell-free virus by centrifugation at 300 g for 5 minutes followed by filtration of the supernatant through a 0.45 micron filter (GVS). 1.2 × 10^6^ infected cells or supernatant from 1.2 × 10^6^ infected cells were added to new uninfected target cells such that the final number of cells in the culture was 6 × 10^6^ at a concentration of 1 × 10^6^cells/ml. EFV at a concentration of 20nM or 40nM was was then added to the cultures. Thereafter, cultures were maintained every 48 hours by the addition of new uninfected target cells at a 1 : 3 ratio of infected to fresh cells. In the experiments where FTC was added, FTC at the concentration of 770nM in addition to the the EFV was added 4 days after addition of 20nM EFV alone. Thereafter, cultures were maintained at a maximum of 1% infected cells by adding fresh cells every 48 hours.

### Sequencing for detection of drug resistant mutants

Genomic DNA from approximately 10^5^ − 10^6^ cells was extracted from the cultures of the evolution experiments every 48 hours using *Quick*-DNA miniprep kits (Zymo Research). The HIV RT gene amplified by PCR from the proviral RNA using Phusion hot start II DNA polymerase (New England Bio-labs) PCR reaction mix. The amplicons were sequenced either by Illumina Miseq or Ion Torrent PGM. Illumina forward primer was 5’-tcgtcggcagcgtcagatgtgtataagagacag TTAATAAGAGAACTCAAGATTTC-3’ and the reverse primer 5’-gtctcgtgggctcggagatgtgtataagagacag CAGCACTATAGGCTGTACTGTC-3’, where lower case sequences are adaptors for Illumina sequencing. Ion Torrent forward primer was 5’-TTAATAAGAGAACTCAAGATTTC-3’ and reverse primer was 5’-CATCTGTTGAGGTGGGGATTTACC-3’. The PCR amplicons were visualized on a 1% agarose gel and bands of the correct size were excised an X-tracta Gel Extraction Tool (Sigma) and product extracted using the QIAquick gel extraction kit (Qiagen). Input DNA for sequencing was quantified using the Qubit dsDNA HS Assay system. For Illumina sequencing, input DNA was diluted in molecular-grade water to reach the starting concentration of 0.2ng/*µ*l. Barcodes were added to each sample using the Nextera XT Index kit (Illumina, Whitehead Scientific, SA). Barcoded amplicons were purified using Ampure XP beads (Beckman Coulter, Atlanta, Georgia) and fragment analysis was performed using the LabChip GX Touch (Perkin Elmer, Waltham, US). The library was pooled at a final concentration of 4nM and further diluted to 3pM. The library was spiked with 20% PhiX plasmid due to the low diversity of the amplicon library. The spiked library was run on a Miseq v2 with a Nano Reagent kit (Illumina). For Ion Torrent sequencing, input DNA was purified using AMPure XP beads (Beckman Coulter, Atlanta, Georgia). Barcoded adapters were added using Ion Plus Fragment Library Kit (Thermo Fisher Scientific) according to manufacturers instructions. Barcoded amplicons were quatified by qPCR using Ion library Quantitation Kit (Thermo Fisher Scientific) and diluted to a concentration of 100pM. Libraries were then pooled and loaded onto chips and sequenced on Ion Torrent PGM using Ion PGM Hi-Q Sequencing Kit (Thermo Fisher Scientific). Fast-q or BAM files from sequencing runs were analysed in Geneious. Drug resistant mutations were found based on a minimum variant frequency of 10^−3^.

## Acknowledgments

This work was supported by National Institute of Allergy and Infectious Diseases, National Institutes of Health Grant 1R01AI138546. JH is supported by a fellowship from the South African National Research Foundation.

---

**SFig. 1.**
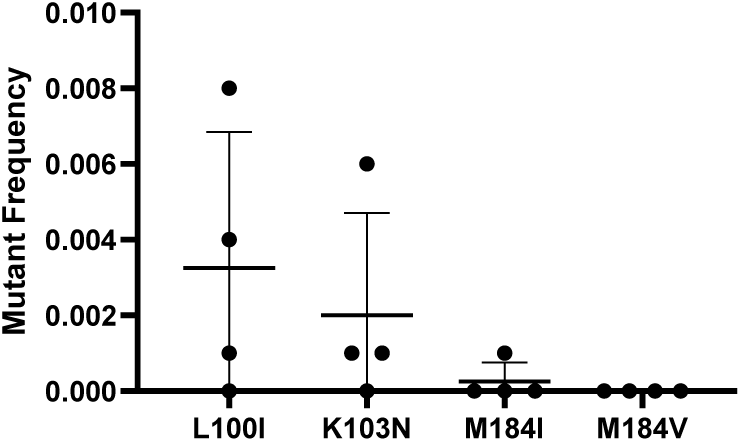
Mutant frequencies measured at start of evolution experiments after two cycles of infection in the absence of drug. The mean frequency for L100I, K103N and M184I were 3 × 10^−2^ ± 4 × 10^−2^, 2 × 10^−2^ ± 3 × 10^−2^, 2 × 10^−3^ ± 5 × 10^−3^ respectively. The mutation frequency for M184V was below the detection threshold of 10^−3^.

**SFig. 2.**
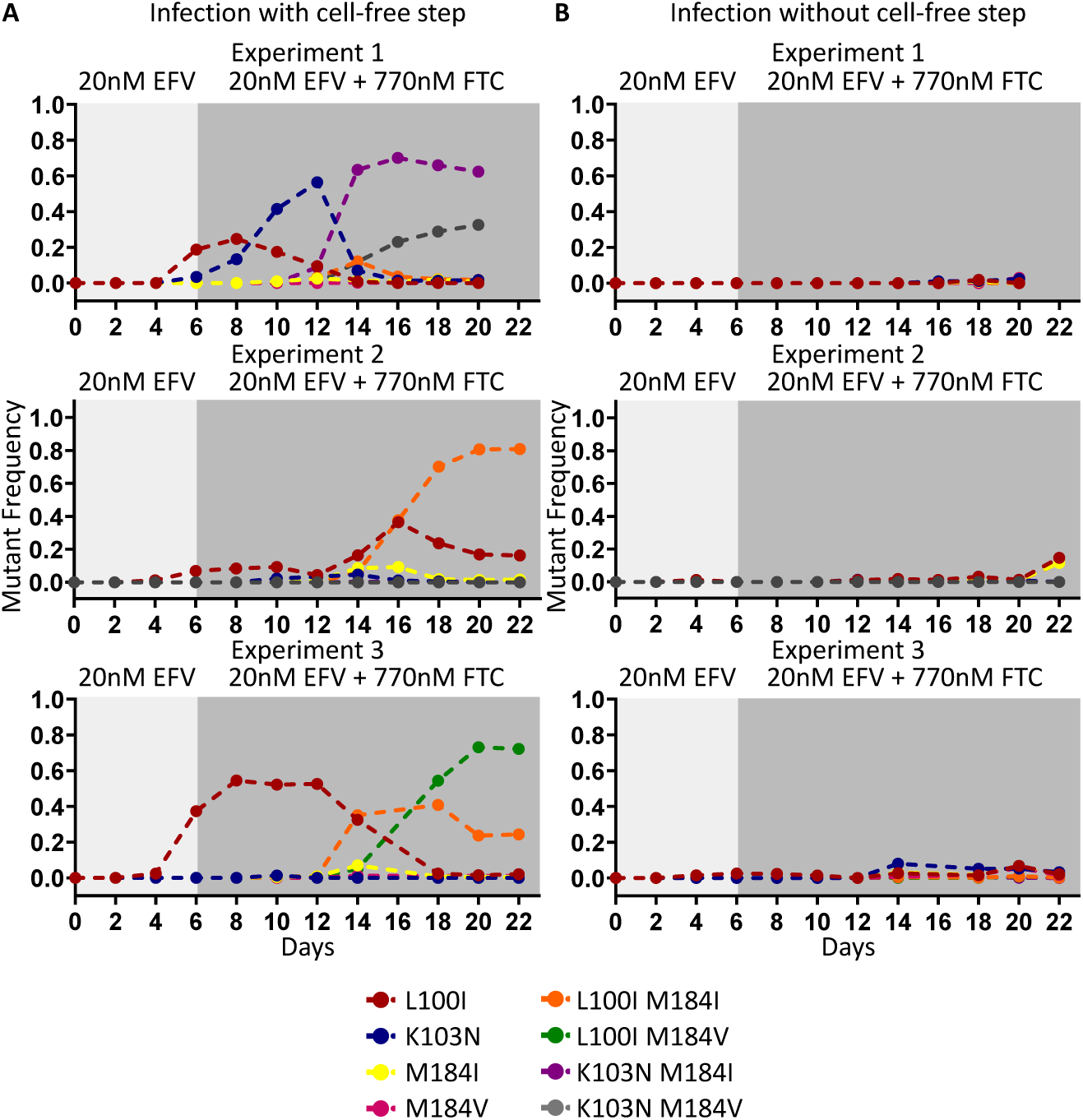
Frequencies of single and linked multidrug resistant mutations. A) Infection in the presence of a cell-free step and B) absence of a cell-free step for 3 experiments. Light grey background shading indicates the days at which the infection was carried out under 20nM EFV monotherapy. Dark grey background shading indicates the days at which the infection in the face of 20nM EFV and 770nM FTC).

